## Supplemental Figures and Tables for "Robust differentiation of human enteroendocrine cells from intestinal stem cells"

Supplementary Figure 1

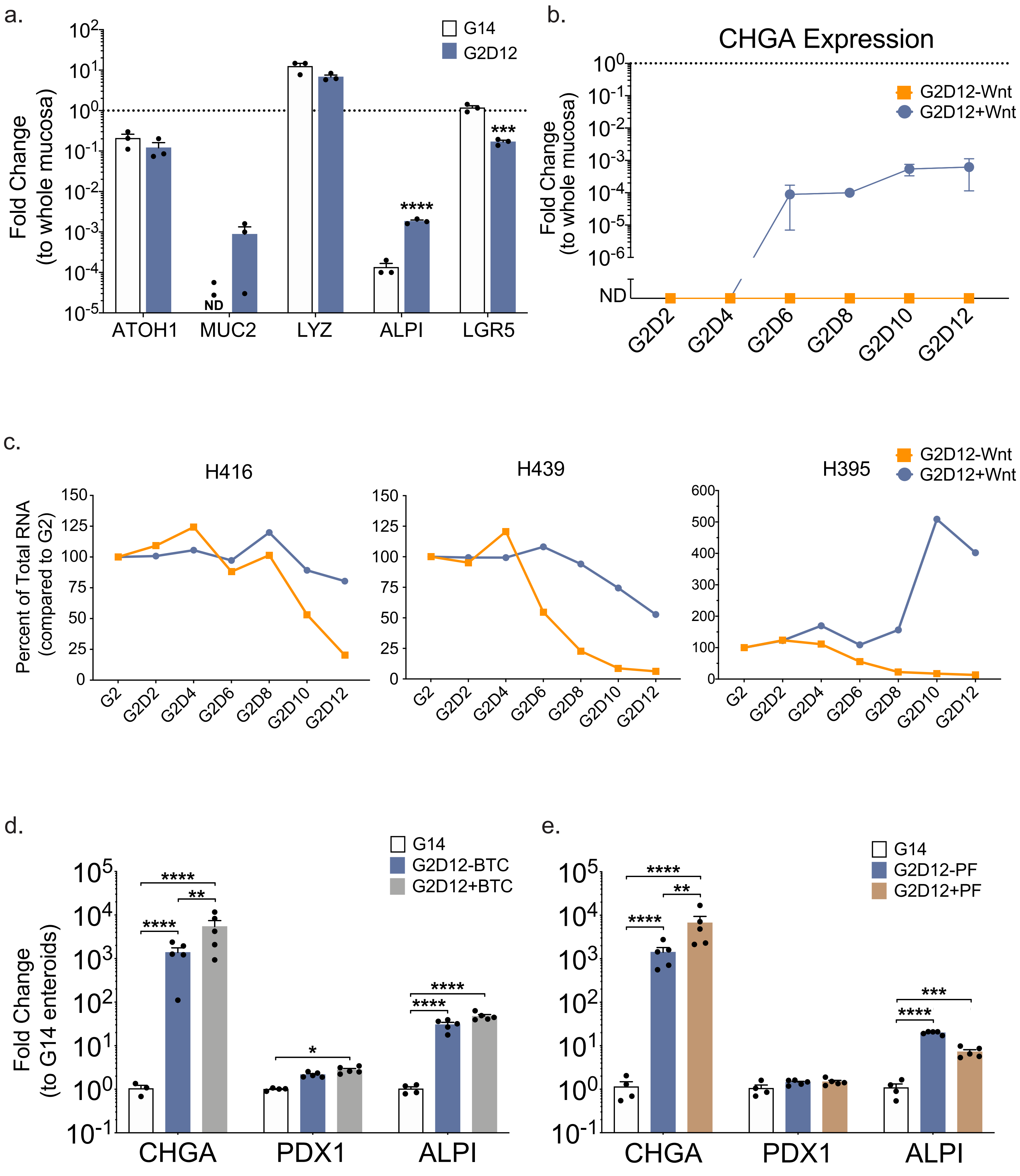

**Supplementary Figure 1. Base Differentiation Media Components Important for Enteroendocrine Marker Expression**

(a) qPCR analysis of intestinal lineage markers of enteroids grown in G14 or G2D12 compared to whole duodenal mucosa and normalized to *18S*. Dotted line denotes expression level in mucosa. Representative experiment showing n = 3 wells from each condition from a single line. *ATOH1* = atonal BHLH transcription factor 1, *MUC2* = mucin 2, *LYZ* = lysozyme, *ALPI* = intestinal alkaline phosphatase, *LGR5* = leucine-rich repeat-containing G-protein coupled receptor 5, ND = not detectable in one or more samples. ***p = 0.004; ****p < 0.0001.

(b) qPCR analysis of chromogranin A (*CHGA*) expression over time of enteroids grown in G2D12 with Wnt (G2D12+Wnt) or G2D12 without Wnt (G2D12-Wnt) compared to whole duodenal mucosa and normalized to *18S*. Dotted line denotes expression level in mucosa. Representative experiment showing n = 3 wells from each condition and timepoint from a single line. For G2D12+Wnt, only two wells expressed *CHGA* at G2D8 with the nondetectable sample being excluded from analysis. ND = not detectable.

(c) Time course study of total RNA levels from three enteroid lines grown in G2D12+Wnt or G2D12-Wnt, shown as a percent compared to RNA levels two days after starting experiment (G2). Representative experiment using an average of n = 3 wells from each condition and timepoint.

(d,e) qPCR analysis of intestinal lineage markers of enteroids grown in (**d**) G2D12 with betacellulin (G2D12+BTC) or G2D12 without betacellulin (G2D12-BTC) and in (**e**) G2D12 with PF06260933 (G2D12+PF) or G2D12 without PF06260933 (G2D12-PF) compared to G14 enteroids and normalized to *18S*. Representative experiments showing n = 3-5 wells from each condition from a single line. *PDX1* = pancreatic and duodenal homeobox 1. (**d**) *p = 0.0478; **p = 0.0023; ****p < 0.0001; (**e**) **p = 0.0080; ***p = 0.0003; ****p < 0.0001.

Bar and line (b) graphs show mean ± SEM; two-way ANOVA with Tukey correction for multiple comparisons (a,d,e). Each experiment repeated with at least three different enteroid lines.

Unless otherwise stated, specific conditions were excluded from statistical analysis if the data from one or more samples was labeled as not detectable. Source data are provided as a Source Data file.

Supplementary Figure 2

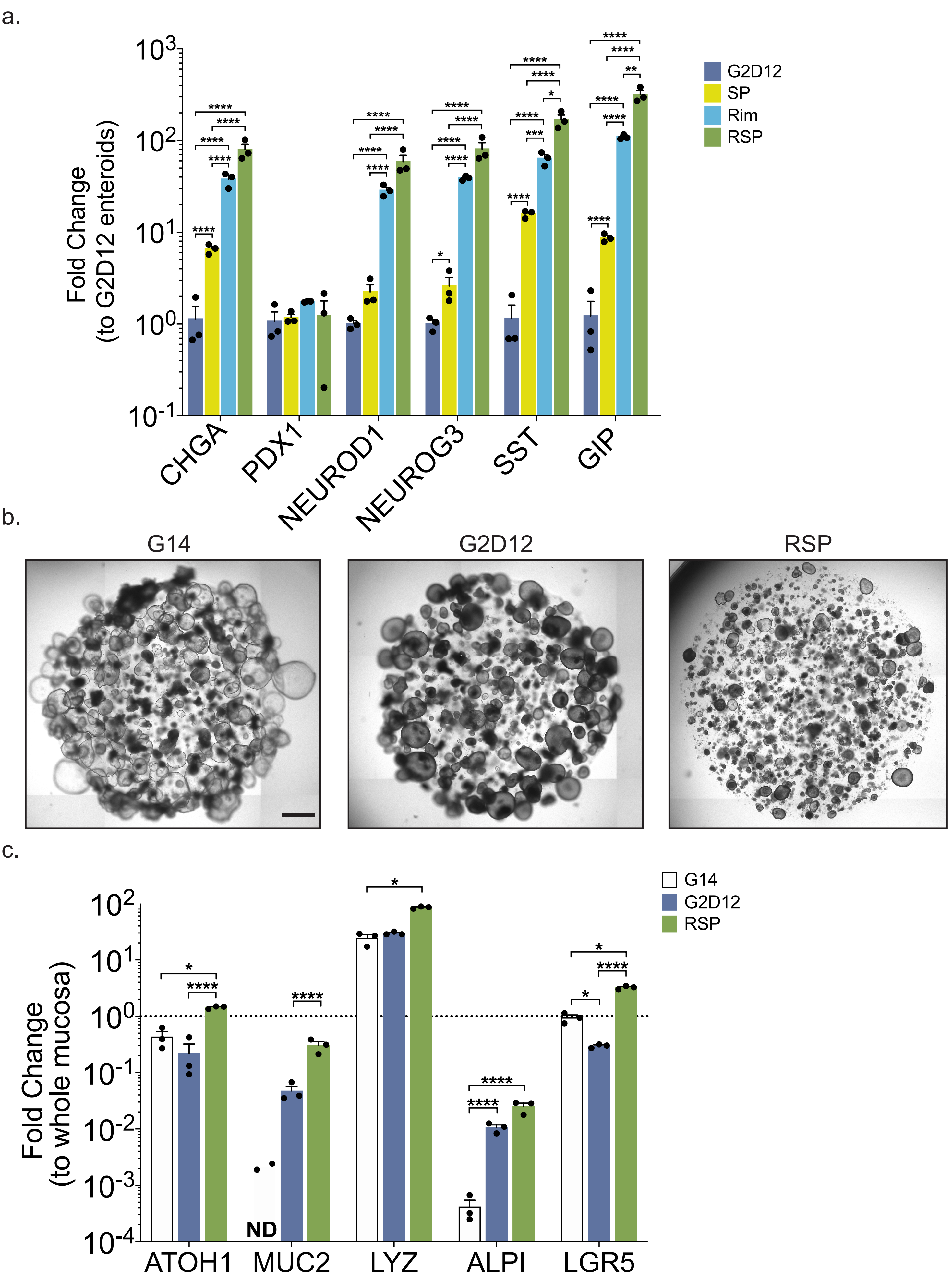

**Supplementary Figure 2. Combination of Rimonabant and SP600125 Induces Enteroendocrine and Other Intestinal Lineage Markers**

(a) qPCR analysis of enteroendocrine markers of enteroids grown in G2D12 with rimonabant (Rim), G2D12 with SP600125 (SP), and G2D12 with rimonabant and SP600125 (RSP) compared to enteroids grown in G2D12 and normalized to *18S*. Representative experiment showing n = 3 wells from each condition from single enteroid line. *CHGA* = chromogranin A, *PDX1* = pancreatic and duodenal homeobox 1, *NEUROD1* = neuronal differentiation 1, *NEUROG3* = neurogenin 3, *SST* = somatostatin, *GIP* = glucose-dependent insulinotropic peptide. *p = 0.0285 (*NEUROG3*), 0.0158 (*SST*); **p = 0.0074; ***p = 0.0003; ****p < 0.0001.

(b) Representative light microscopy of enteroids (whole well) grown in G14, G2D12, and RSP. Scale bar = 1mm.

(c) qPCR analysis of intestinal lineage markers of enteroids grown in G14, G2D12, and RSP compared to whole duodenal mucosa and normalized to *18S*. Dotted line denotes expression level in whole duodenal mucosa. Representative experiment showing n = 3 wells from each condition from a single enteroid line. *ATOH1* = atonal BHLH transcription factor 1, *MUC2* = mucin 2, *LYZ* = lysozyme, *ALPI* = intestinal alkaline phosphatase, *LGR5* = leucine-rich repeat-containing G-protein coupled receptor 5, ND = not detectable in one or more samples. *p = 0.0118 (*ATOH1*), 0.0108 (*LYZ*), 0.0334 (*LGR5*, G14 to G2D12), 0.0133 (*LGR5*, G14 to RSP); ****p < 0.0001.

Bars show mean ± SEM; two-way ANOVA with Tukey correction for multiple comparisons (a,c). Each experiment repeated with at least three different enteroid lines. Specific conditions were excluded from statistical analysis if the data from one or more samples was labeled as not detectable. Source data are provided as a Source Data file.

Supplementary Figure 3

a.

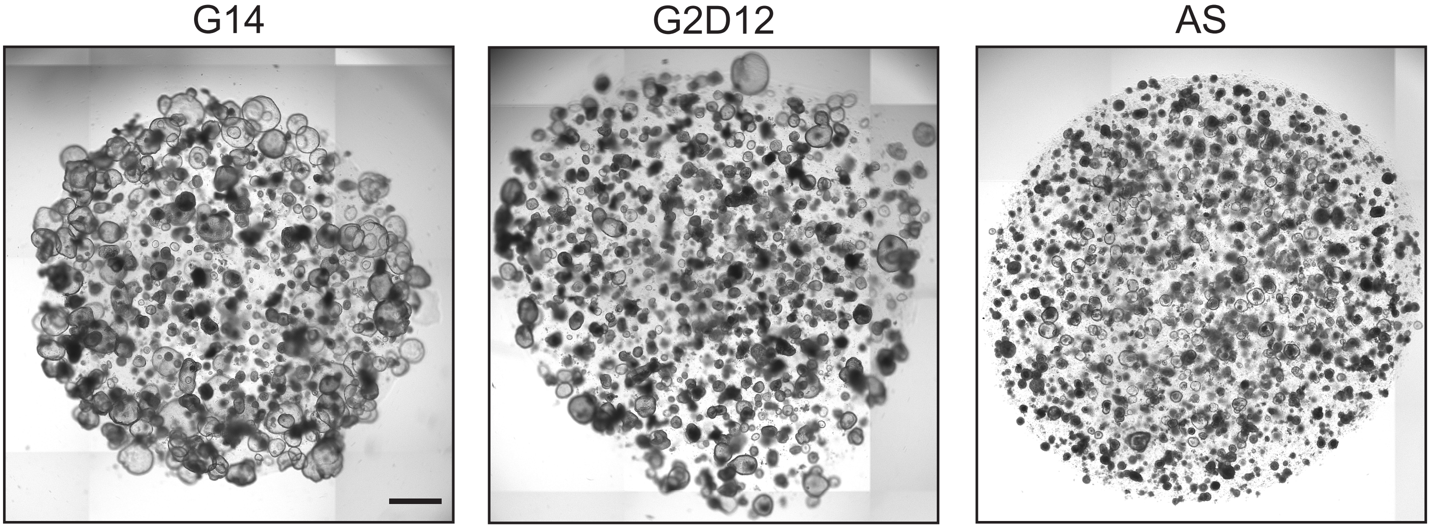


b.

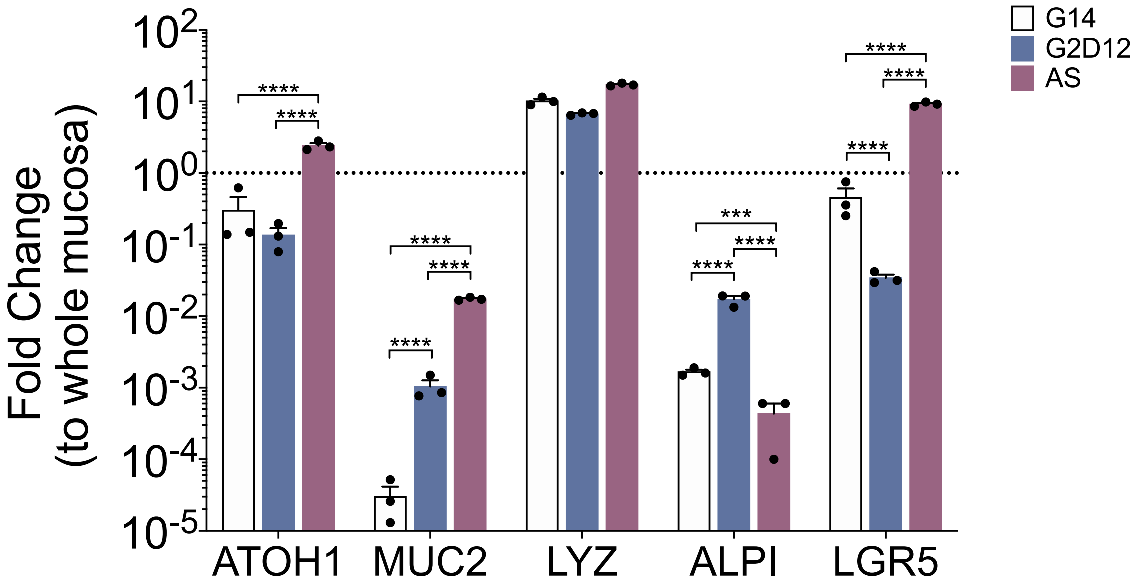

**Supplementary Figure 3. AS1842856 Induces Specific Intestinal Lineage Markers**

(a) Representative light microscopy of enteroids (whole well) grown in G14, G2D12, and G2D12 with AS1842856 (AS). Scale bar = 1mm.

(b) qPCR analysis of intestinal lineage markers of enteroids grown in G14, G2D12, and AS compared to whole duodenal mucosa and normalized to *18S*. Dotted line denotes expression level in whole duodenal mucosa. Representative experiment showing n = 3 wells from each condition from single enteroid line. *ATOH1* = atonal BHLH transcription factor 1, *MUC2* = mucin 2, *LYZ* = lysozyme, *ALPI* = intestinal alkaline phosphatase, *LGR5* = leucine-rich repeat-containing G-protein coupled receptor 5. ***p = 0.001, ****p < 0.0001.

Bars show mean ± SEM; two-way ANOVA with Tukey correction for multiple comparisons. Each experiment repeated with at least three different enteroid lines. Source data are provided as a Source Data file.

Supplementary Figure 4

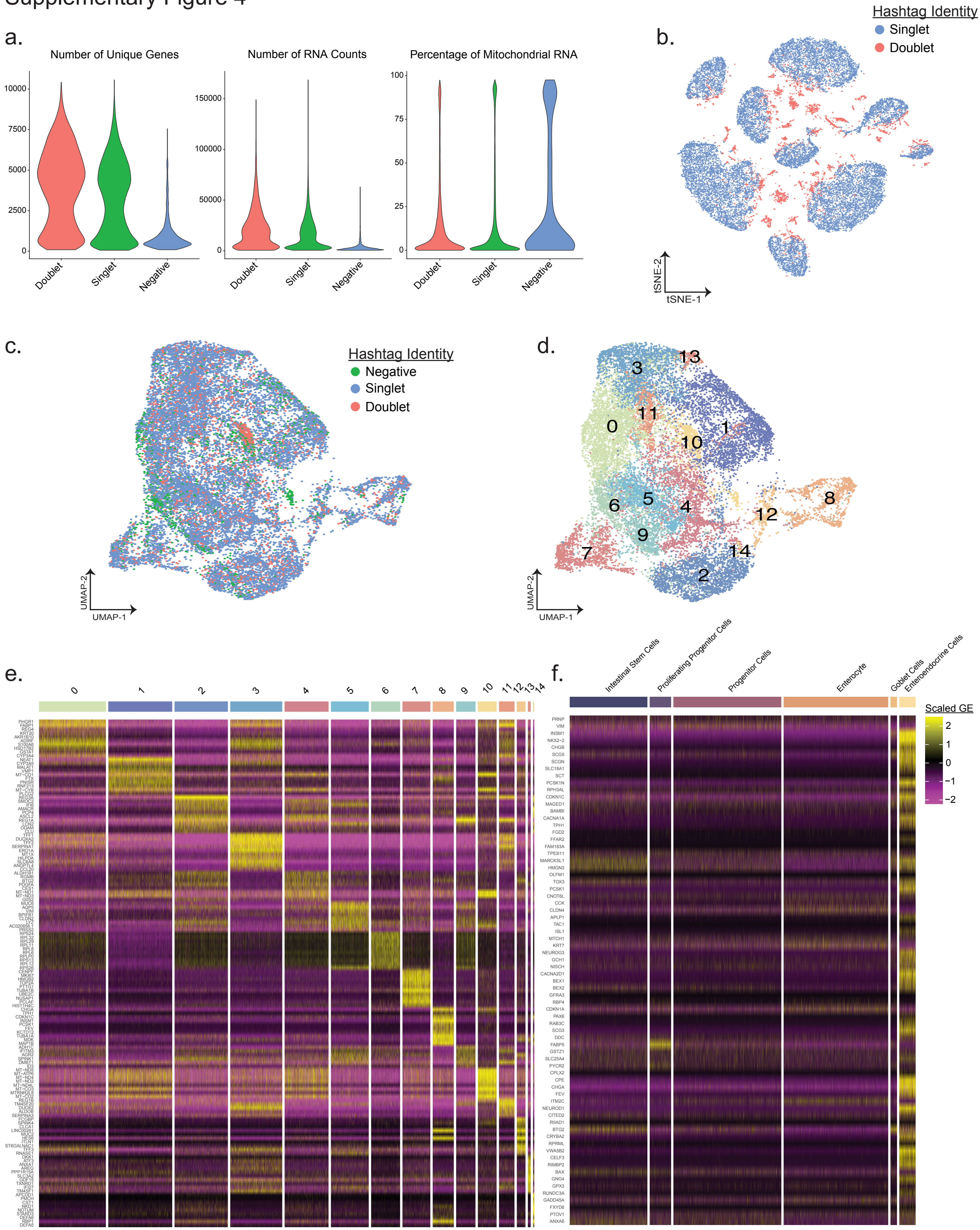

**Supplementary Figure 4. Cellular Validation and Characterization of Enteroid Single Cell Transcriptomes**

(a) Violin plots summarizing the distribution of genes (left), RNA counts or unique molecular identifiers (center), and the proportion of mitochondrial genes (right) within the scRNA-seq dataset.

(b) t-distributed stochastic neighbor embedding (tSNE) visualization of 23,335 cells from all samples with color denoting the hashtag identity. Singlet cells were denoted as having significant enrichment of only one hashtag signal per individual cell. Doublet cells had significant enrichment of more than one hashtag signal.

(c) Uniform manifold approximation and projection (UMAP) visualization of 25,673 cells originating from all samples, with color being used to distinguish between negative, doublet, or singlet cells.

(d) UMAP visualization from (c) with cells labeled by their Louvain cluster identity.

(e) Heatmap of the top 10 most differentially expressed genes per Louvain cluster compared to all other clusters. Differentially expressed genes were calculated using the Wilcoxon rank sum test and the top 10 genes of each cluster were selected based on their log fold enrichment in the specified cluster compared to all other clusters. Heatmap values corresponded to row-scaled Pearson residuals of gene expression that was normalized using regularized negative binomial regression.

(f) Heatmap of marker genes significantly enriched in *in vivo* EE cells isolated from the mouse small intestinal epithelial cell atlas[1]. Heatmap values for display were calculated using similar methods seen in (e) and may not represent significantly enriched genes.

Supplementary Figure 5

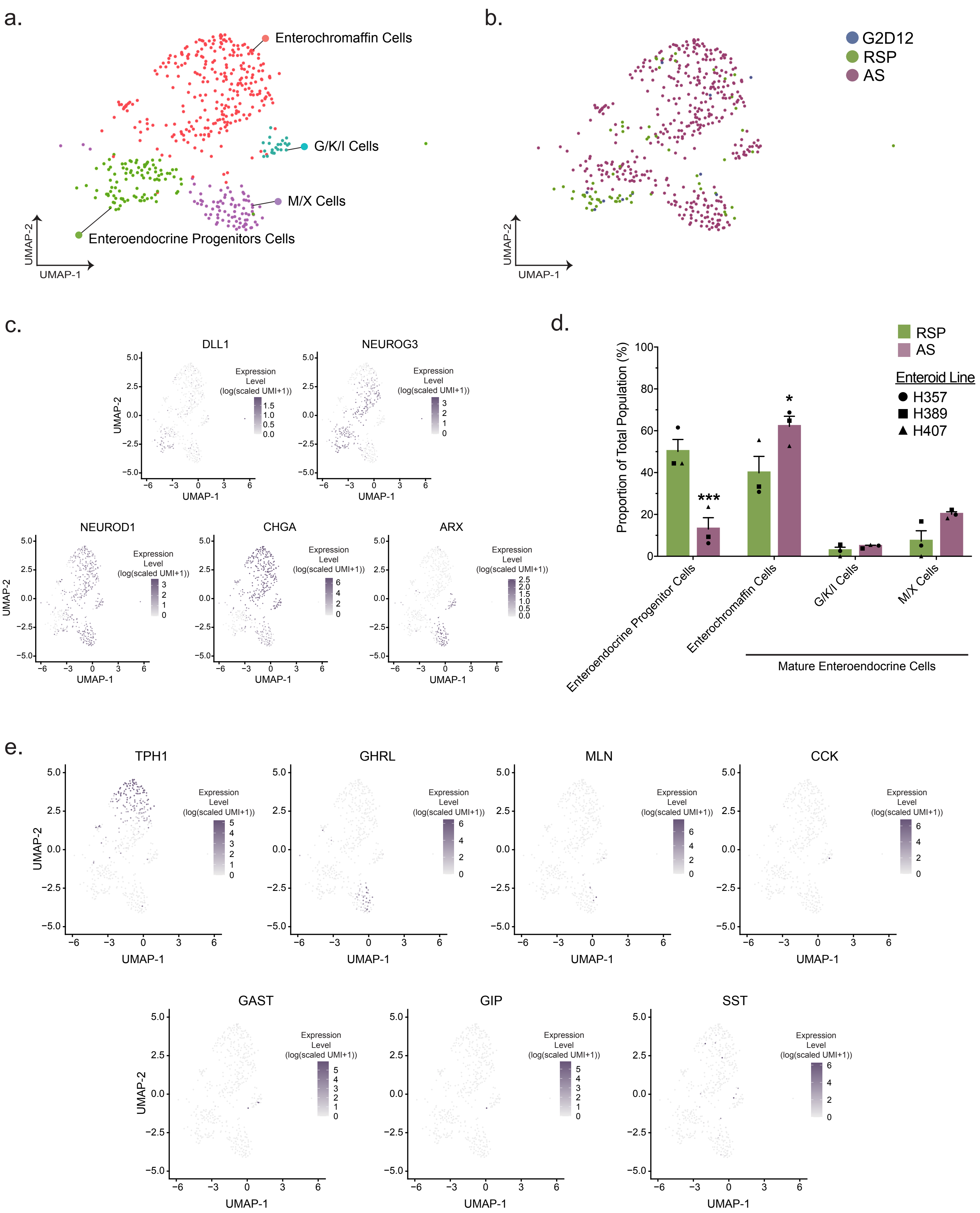

**Supplementary Figure 5. Single Cell Analysis of Enteroendocrine Cells**

(a) Uniform manifold approximation and projection (UMAP) visualization of 471 enteroendocrine cells from all samples and conditions, colored by cell identity.

(b) UMAP visualization from (a), colored by enteroid differentiation condition.

(c) Marker gene overlay for binned count-based expression level (log(scaled UMI + 1)) of various enteroendocrine marker genes, projected on the UMAP from (a). *DLL1* = delta like canonical notch ligand 1, *NEUROG3* = neurogenin 3, *NEUROD1* = neuronal differentiation 1, *CHGA* = chromogranin A, *ARX* = aristaless-related homeobox.

(d) Proportional abundance of enteroendocrine subsets by culture condition. G2D12 was not included due to low number of cells. Each culture condition consists of three different enteroid lines from distinct human donors, as denoted by data point shape. Bars show mean ± SEM; two-way ANOVA with Tukey correction for multiple comparisons. *p = 0.0160; ***p < 0.0002.

(e) Marker gene overlay for binned count-based expression level (log(scaled UMI + 1)) of various enteroendocrine hormone genes, projected on the UMAP from (a). *TPH1* = tryptophan hydroxylase 1, *GHRL* = ghrelin, *MLN* = motilin, *CCK* = cholecystokinin, *GAST* = gastrin, *GIP* = glucose-dependent insulinotropic peptide, *SST* = somatostatin. Source data are provided as a Source Data file.

Supplementary Figure 6

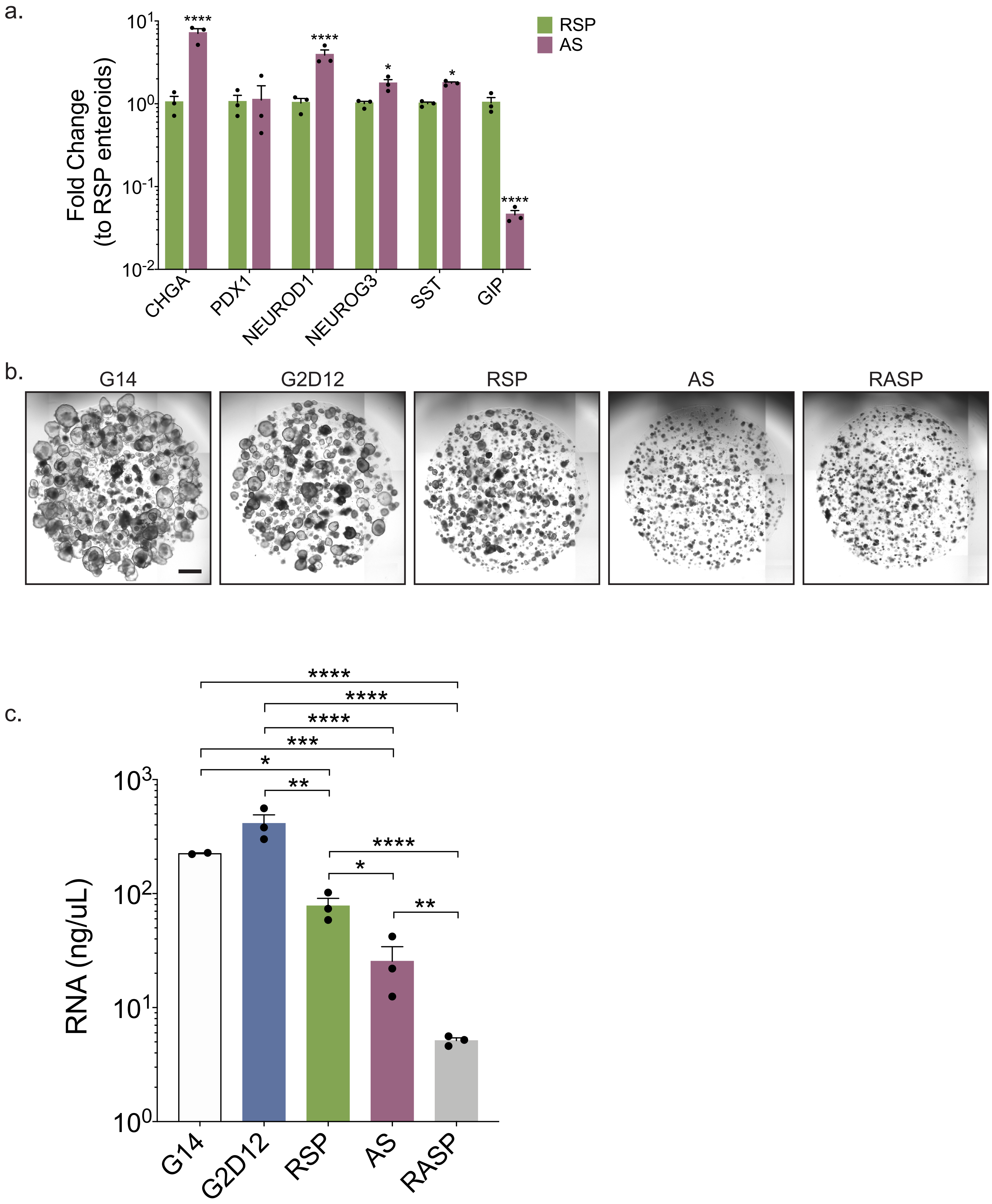

**Supplementary Figure 6. Combination of AS1842856 and Rimonabant/SP600125 For All of Differentiation Leads to Reduction in Isolated RNA**

(a) qPCR analysis of enteroendocrine markers of enteroids grown in AS compared to RSP and normalized to *18S*. Representative experiment showing n = 3 wells from each condition from a single enteroid line. *CHGA* = chromogranin A, *PDX1* = pancreatic and duodenal homeobox 1, *NEUROD1* = neuronal differentiation 1, *NEUROG3* = neurogenin 3, *SST* = somatostatin, *GIP* = glucose-dependent insulinotropic peptide. *p = 0.0407 (*NEUROG3*), 0.0306 (*SST*); ****p < 0.0001.

(b) Representative light microscopy of enteroids (whole well) grown in G14, G2D12, RSP, AS, and G2D12 with AS and RSP (RASP). Scale bar = 1 mm.

(c) Total RNA levels from enteroids grown in G14, G2D12, RSP, AS, and RASP. Representative results from n = 2-3 wells from each condition from a single enteroid line. *p = 0.0487 (G14 to RSP), 0.0142 (AS to RSP); **p = 0.0018 (G2D12 to RSP), 0.0039 (AS to RASP); ***p = 0.0004; ****p < 0.0001.

Bars show mean ± SEM; two-way ANOVA with Tukey correction for multiple comparisons (a); one-way ANOVA with Tukey correction for multiple comparisons (c). Each experiment repeated with at least three different enteroid lines. Source data are provided as a Source Data file.

Supplementary Figure 7

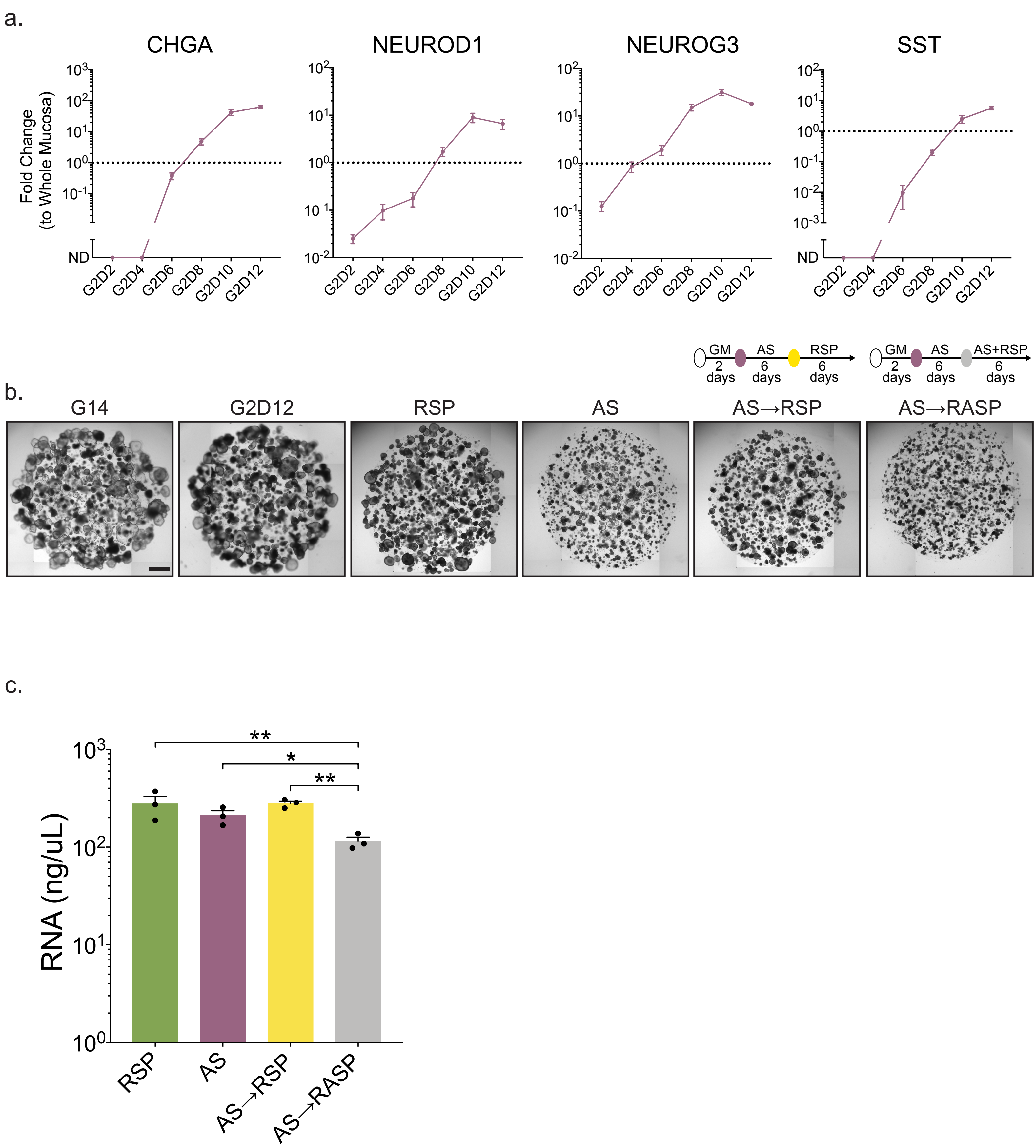

**Supplementary Figure 7. Combination of AS1842856 and Rimonabant/SP600125 After Initial AS1842856 Exposure Yields Viable Enteroids**

(a) qPCR analysis of enteroendocrine marker expression over time from enteroids grown in AS compared to whole duodenal mucosa and normalized to *18S*. RNA was collected every two days after start of differentiation. Dotted line denotes expression level in whole duodenal mucosa. Representative experiment showing n = 3 wells from each timepoint, except n = 2 for G2D12, from a single enteroid line. At G2D2, only two of three wells expressed *NEUROD1* and *NEUROG3*, with nondetectable samples excluded from analysis. *CHGA* = chromogranin A, *NEUROD1* = neuronal differentiation 1, *NEUROG3* = neurogenin 3, *SST* = somatostatin, ND = not detectable.

(b) Representative light microscopy of enteroids (whole well) grown in G14, G2D12, RSP, AS, AS→RSP, and AS→RASP. Specific culture schematics of AS→RSP and AS→RASP located above each panel, respectively. Scale bar = 1 mm.

(c) Total mRNA levels from enteroids grown in RSP, AS, AS→RSP, and AS→RASP. Representative experiment showing n = 3 wells from each condition from a single enteroid line. *p = 0.0456; **p = 0.0071 (RSP to AS→RASP), 0.0052 (AS→RSP to AS→RASP).

Bars and line graph show mean ± SEM; one-way ANOVA with Tukey correction for multiple comparisons (c). Each experiment repeated with at least three different enteroid lines. Source data are provided as a Source Data file.

Supplementary Figure 8

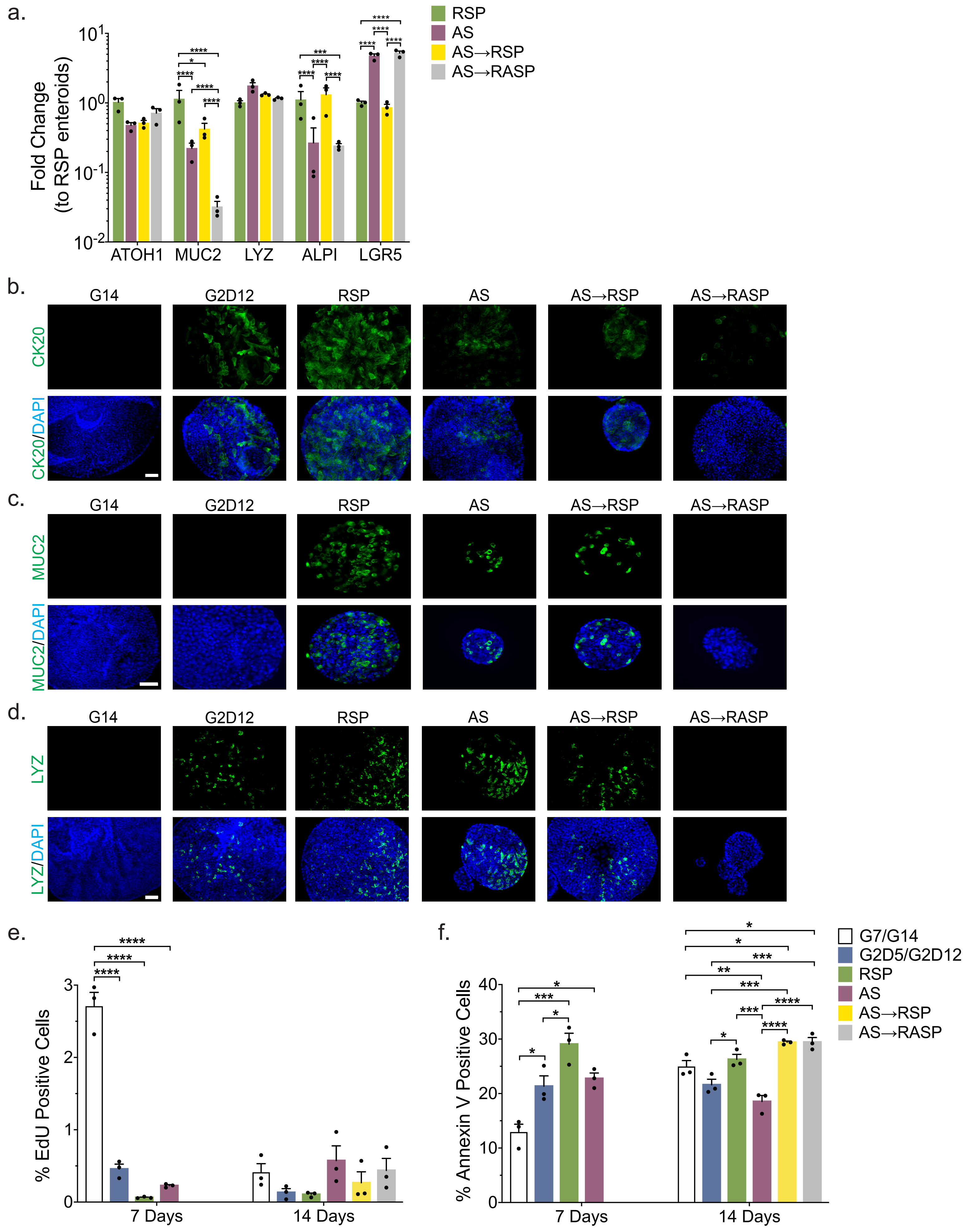

**Supplementary Figure 8. Induction of Specific Intestinal Lineage Markers, Cell Viability, and Proliferation Compared Across Differentiation Methods**

(a) qPCR analysis of intestinal lineage markers of enteroids grown in AS, AS→RSP, and AS→RASP compared to RSP and normalized to *18S*. Representative experiment showing n = 3 wells from each condition from single enteroid line. *ATOH1* = atonal BHLH transcription factor 1, *MUC2* = mucin 2, *LYZ* = lysozyme, ALPI = intestinal alkaline phosphatase, *LGR5* = leucine-rich repeat-containing G-protein coupled receptor 5. *p = 0.0266; ***p < 0.0002; ****p < 0.0001.

(b) Representative immunofluorescence staining of cytokeratin 20 (CK20, green) in enteroids treated with G14, G2D12, RSP, AS, AS→RSP, and AS→RASP. DNA (4′,6-diamidino-2-phenylindole (DAPI), blue). Scale bar = 50 µm.

(c) Representative immunofluorescence staining of mucin 2 (MUC2, green) in enteroids treated with G14, G2D12, RSP, AS, AS→RSP, and AS→RASP. DNA (4′,6-diamidino-2-phenylindole (DAPI), blue). Scale bar = 50 µm.

(d) Representative immunofluorescence staining of lysozyme (LYZ, green) in enteroids treated with G14, G2D12, RSP, AS, AS→RSP, and AS→RASP. DNA (4′,6-diamidino-2-phenylindole (DAPI), blue). Scale bar = 50 µm.

(e) Percentage of EdU-positive cells after a two-hour chase with 10 µM EdU in enteroids grown in their respective medias at either 7 days (left side) or 14 days (right side). ****p < 0.0001.

(f) Percentage of Annexin V-positive cells in enteroids grown in their respective medias at either 7 days (left side) or 14 days (right side). *p = 0.0276 (G7 to G2D5), 0.0121 (G7 to AS), 0.0428 (G2D5 to RSP), 0.0459 (G14 to AS→RSP), 0.0440 (G14 to AS→RASP), 0.0371 (G2D12 to RSP); **p = 0.0043; ***p = 0.0005 (G7 to RSP), 0.0008 (G2D12 to AS→RSP), 0.0007 (G2D12 to AS→RASP), 0.0008 (RSP to AS); ****p < 0.0001.

Bars and line graph show mean ± SEM; two-way ANOVA with Tukey correction for multiple comparisons (a); one-way ANOVA with Tukey correction for multiple comparisons (e,f) as comparisons between 7 and 14 days were not made. Each experiment repeated with at least three different enteroid lines. Source data are provided as a Source Data file.

Supplementary Figure 9

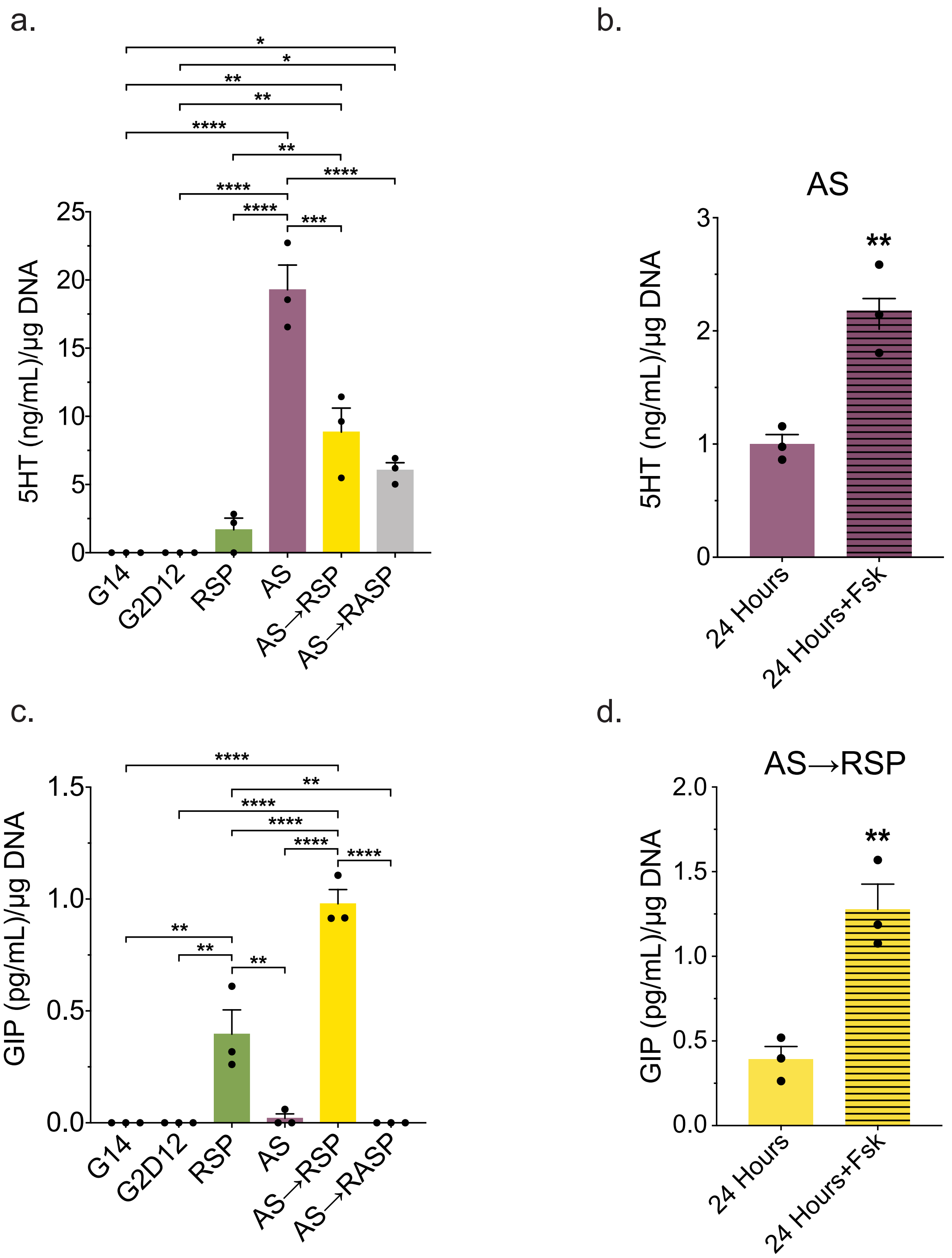

**Supplementary Figure 9. Hormone Secretion of Different Conditions Controlled for Total DNA**

(a) Serotonin (5HT) ELISA of conditioned media from the last two days of differentiation of enteroids grown in G14, G2D12, RSP, AS, AS→RSP, and AS→RASP controlled for total DNA from each sample. Representative experiment showing n = 3 wells from each condition from a single enteroid line. *p = 0.0223 (G14 to AS→RASP, G2D12 to AS→RASP); **p = 0.0012 (G14 to AS→RSP, G2D12 to AS→RSP), 0.0067 (RSP to AS→RSP); ***p = 0.0003, ****p < 0.0001.

(b) 5HT ELISA of AS conditioned media collected after 24 hours on day 13 (solid bar) and after 24 hours with forskolin (Fsk) on day 14 (striped bar) controlled for total DNA from each sample. Representative experiment showing n = 3 wells from each condition from a single enteroid line. **p = 0.0081.

(c) Glucose-dependent insulinotropic peptide (GIP) ELISA of conditioned media from the last two days of differentiation of enteroids grown in G14, G2D12, RSP, AS, AS→RSP, and AS→RASP controlled for total DNA from each sample. Representative experiment showing n = 3 wells from each condition from a single enteroid line. **p = 0.0017 (G14 to RSP, G2D12 to RSP, RSP to AS→RASP), 0.0027 (AS to RSP); ****p < 0.0001.

(d) GIP ELISA of AS→RSP conditioned media collected after 24 hours on day 13 (solid bar) and after 24 hours with Fsk on day 14 (striped bar) controlled for total DNA from each sample. Representative experiment showing n = 3 wells from each condition from a single enteroid line. **p = 0.0061.

Bars show mean ± SEM; one-way ANOVA with Tukey correction for multiple comparisons (a,c); two-tailed unpaired *t* test (b,d). To control for enteroid number, protein concentrations were divided by DNA concentration. Each experiment repeated with at least three different enteroid lines. Source data are provided as a Source Data file.

Supplementary Figure 10

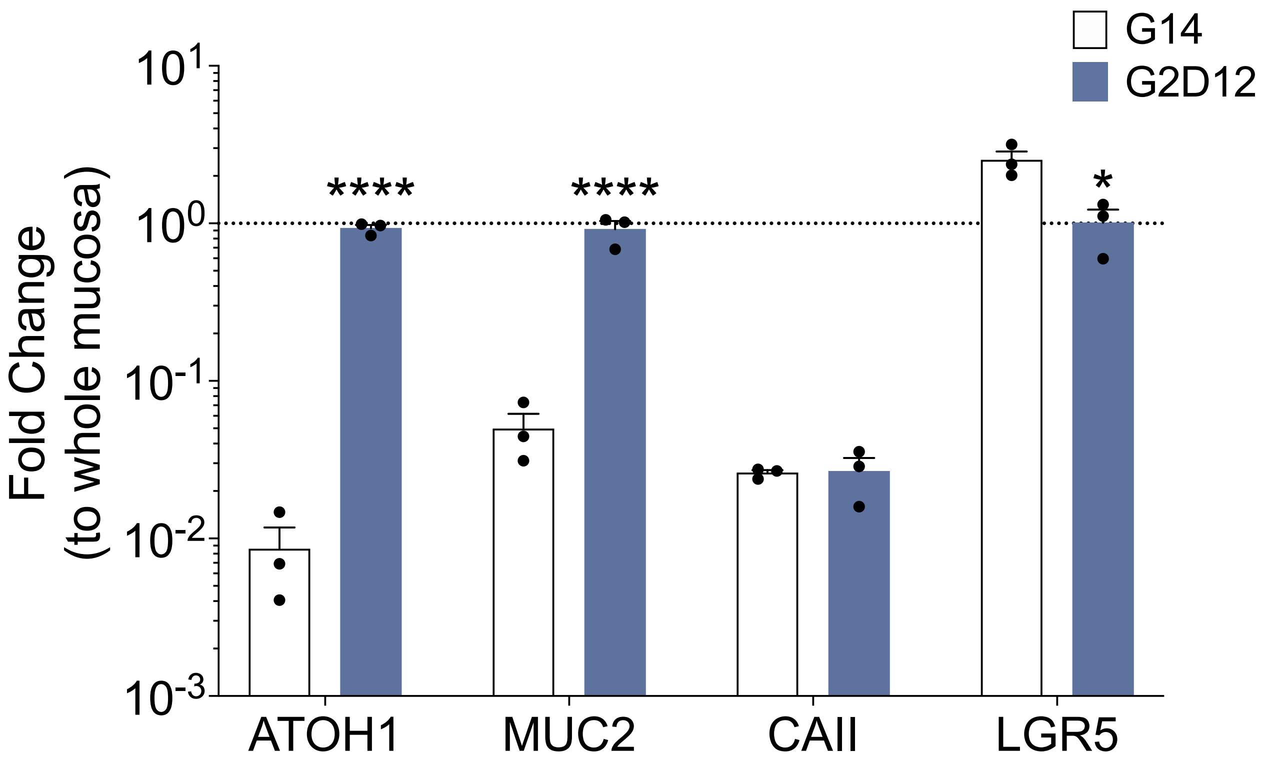

**Supplementary Figure 10. Induction of Other Lineage Markers in Rectoids**

qPCR analysis of intestinal lineage markers of rectoids grown in G14 and G2D12 compared to whole rectal mucosa and normalized to *18S*. Dotted line denotes expression level in whole rectal mucosa. Representative experiment showing n = 3 wells from each condition from a single rectoid line. *ATOH1* = atonal BHLH transcription factor 1, *MUC2* = mucin 2, *CAII* = carbonic anhydrase II, *LGR5* = leucine-rich repeat-containing G-protein coupled receptor 5. *p < 0.0150; ****p < 0.0001.

Bars show mean ± SEM; two-way ANOVA with Tukey correction for multiple comparisons. Experiment repeated with at least three different rectoid lines. Source data are provided as a Source Data file.

Supplementary Figure 11

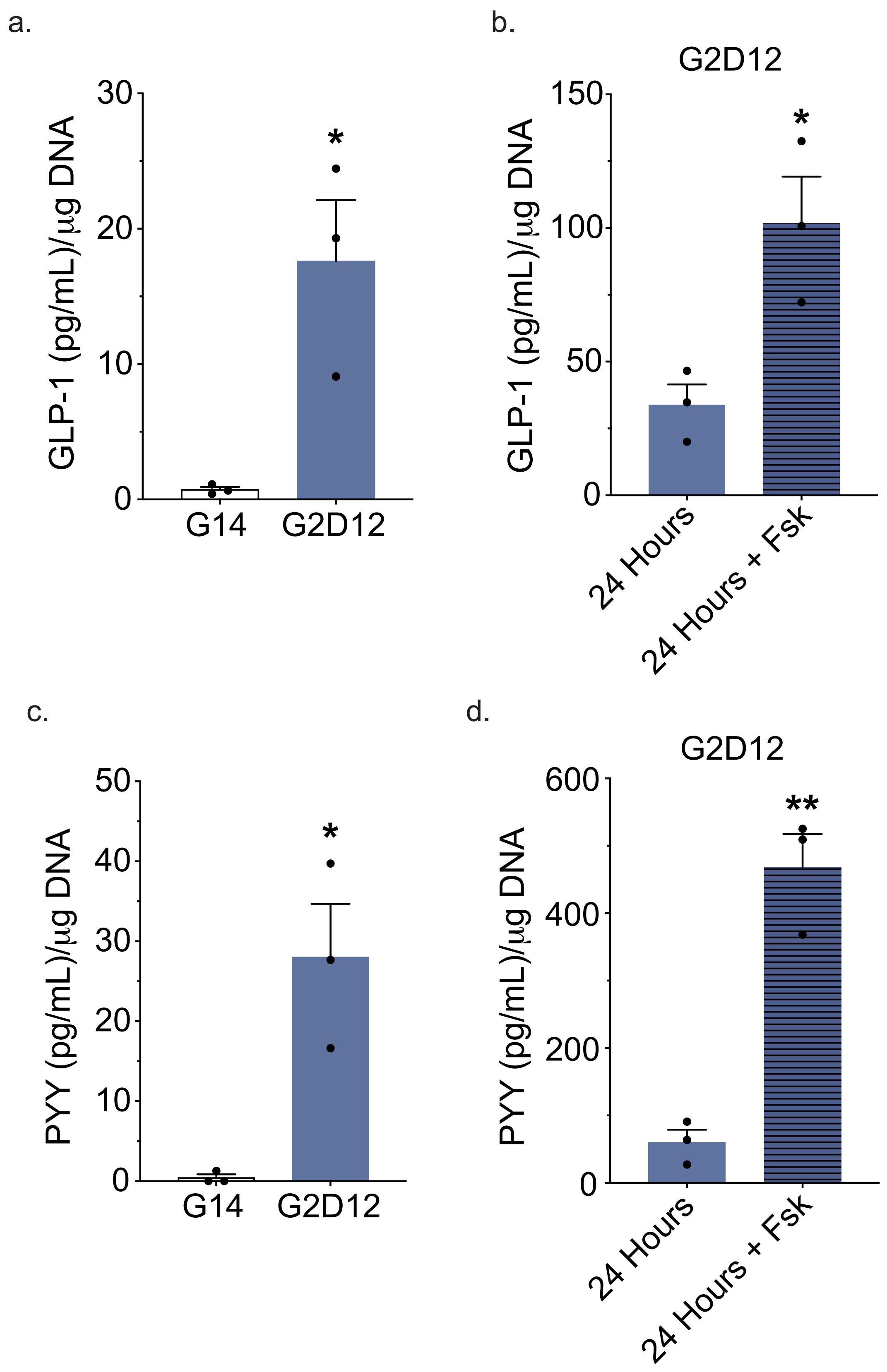

**Supplementary Figure 11. Rectoid Hormone Secretion Controlled for Total DNA**

(a) Glucagon-like peptide-1 (GLP-1) ELISA of conditioned media from the last two days of differentiation of enteroids grown in G14 and G2D12 controlled for total DNA from each sample. Representative experiment showing n = 3 wells from each condition from a single rectoid line. *$p < 0.0202$.

(b) GLP-1 ELISA of G2D12 conditioned media collected after 24 hours on day 13 (solid bar) and after 24 hours with forskolin (Fsk) on day 14 (striped bar) controlled for total DNA from each sample. Representative experiment showing n = 3 wells from each condition from a single rectoid line. *$p < 0.0233$.

(c) Peptide YY (PYY) ELISA of conditioned media from the last two days of differentiation of enteroids grown in G14 and G2D12 controlled for total DNA from each sample. Representative experiment showing n = 3 wells from each condition from a single rectoid line. *$p = 0.0145$.

(d) PYY ELISA of G2D12 conditioned media collected after 24 hours on day 13 (solid bar) and after 24 hours with Fsk on day 14 (striped bar) controlled for total DNA from each sample. Representative experiment showing n = 3 wells from each condition from a single enteroid line. **$p = 0.0016$.

Bars show mean ± SEM; two-tailed unpaired *t* test (a,b,c,d). To control for rectoid number, protein concentrations were divided by DNA concentration. Each experiment repeated with at least three different rectoid lines. Source data are provided as a Source Data file.

Supplementary Figure 12

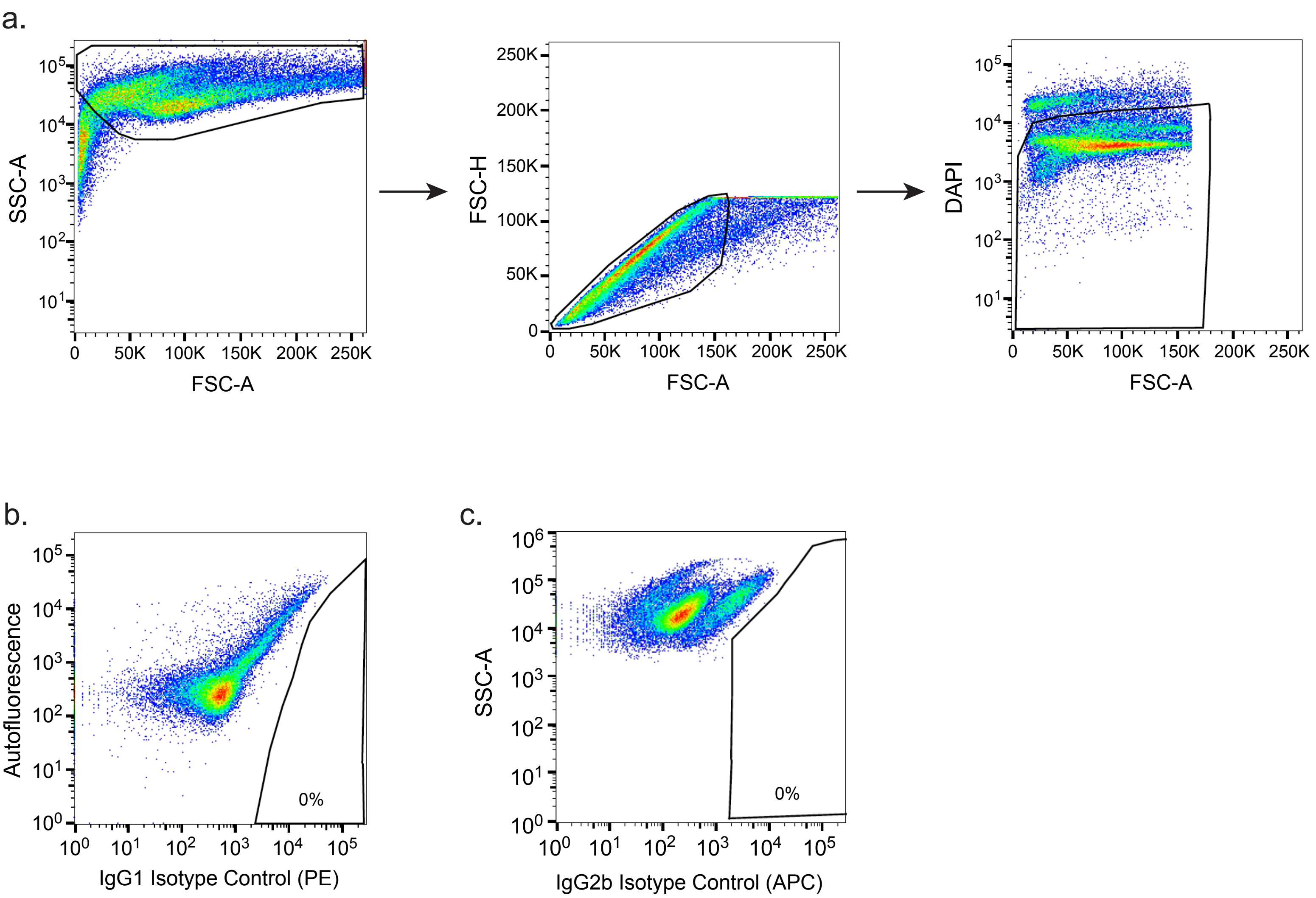

**Supplementary Figure 12. Gating Strategy for Flow Cytometry**

(a) Enteroid cells are differentiated from cellular debris based on their forward and side scatter area (FSC-A and SSC-A, respectively) parameters. Cells are then examined based on their FSC-A and FSC-Height (H) to exclude doublets. 4′,6-diamidino-2-phenylindole (DAPI) staining is then utilized to identify dead cells, with DAPI high-positive cells being excluded from further gating.

(b) The CHGA-positive gate is set by using an IgG1 K isotype control conjugate with phycoerythrin (PE).

(c) The CHGA-positive gate is set by using an IgG2b K isotype control conjugate with Alexa fluor 647 (APC).

Supplementary Table 1. Growth and Differentiation Media Components

| ENTEROID | | RECTOID | |
|---|---|---|---|
| **Growth Media** | **Differentiation Media** | **Growth Media** | **Differentiation Media** |
| L-WRN conditioned media (50% v/v) | | L-WRN conditioned media (65% v/v) | |
| DMEM/F12 (45% v/v) | | DMEM/F12 (30% v/v) | |
| Glutamax (1% v/v) | | Glutamax (1% v/v) | |
| N-2 Supplement (1% v/v) | | N-2 Supplement (1% v/v) | |
| B-27 Supplement (1% v/v) | | B-27 Supplement (1% v/v) | |
| HEPES (10mM) | | HEPES (10mM) | |
| Primocin (100μg/mL) | | Primocin (100μg/mL) | |
| Normocin (100μg/mL) | | Normocin (100μg/mL) | |
| A83-01 (500nM) | | A83-01 (500nM) | |
| N-Acetyl-cysteine (500μM) | | N-Acetyl-cysteine (500μM) | |
| Recombinant Murine EGF (50ng/mL) | | Recombinant Murine EGF (50ng/mL) | |
| Human [Leu15] Gastrin I (50nM) | | Human [Leu15] Gastrin I (50nM) | |
| Nicotinamide (10mM) | DAPT (20μM) | Nicotinamide (10mM) | DAPT (20μM) |
| SB202190 (10μM) | Betacellulin (20ng/mL) | SB202190 (10μM) | Betacellulin (20ng/mL) |
| | Tubastatin-A (10μM) | Prostaglandin-E2 (10nM) | Tubastatin-A (10μM) |
| | PF06260933 (6μM) | | PF06260933 (6μM) |
| | Tranylcypromine (1.5μM) | | Tranylcypromine (1.5μM) |
| | Rimonabant (10μM) | | |
| | SP600125 (10μM) | | |
| | AS1842856 (100nM) | | |

**Supplementary Table 1. Growth and Differentiation Media Components**

Growth factors, supplements, and small molecules common to both growth and differentiation medias are listed first. L-WRN cell line used to produce conditioned media containing Wnt3a, Noggin and R-spondin 3[2]. HA-R-Spondin1-Fc 293T cell line used to produce conditioned media with only R-spondin 1[3]. This conditioned media, with supplemented Noggin (100 ng/mL), was used to make Wnt3a-free differentiation media.

Supplementary Table 2. Description of Samples

| Application | Age Range (years) | Number of Males | Number of Females |
|---|---|---|---|
| Enteroid Lines | 13-21 | 4 | 4 |
| Rectoid Lines | 12-18 | 2 | 6 |
| Duodenal mucosa RNA | 55-82 | 1 | 2 |
| Rectal mucosa RNA | 15-18 | 0 | 3 |

**Supplementary Table 2. Description of Samples**

Samples used to produce organoid cell lines are labeled as either enteroid or rectoid lines, while those used as a duodenal or rectal whole mucosa control are labeled as mucosa RNA. For each application, the age range and number of males and females are noted. All biopsies and resections were determined to be healthy, and from patients without known gastrointestinal disease, before being provided to the researchers de-identified, with age and sex not known until after completion of experiments.

Supplementary Table 3. Taqman qPCR Primers

| Name | Abbreviation | Identifier |
|---|---|---|
| 18S | *18S* | Hs99999901_s1 |
| Intestinal alkaline phosphatase | *ALPI* | Hs00357579_g1 |
| Atonal bHLH transcription factor 1 | *ATOH1* | Hs00944192_s1 |
| Carbonic anhydrase II | *CAII* | Hs01070108_m1 |
| Cholecystokinin | *CCK* | Hs00174937_m1 |
| Chromogranin A | *CHGA* | Hs00900370_m1 |
| GATA binding protein 4 | *GATA4* | Hs00171403_m1 |
| Glucose-dependent insulinotropic polypeptide | *GIP* | Hs00175030_m1 |
| Glucagon | *GCG* | Hs01031536_m1 |
| Leucine-rich repeat-containing G-protein coupled receptor 5 | *LGR5* | Hs00969422_m1 |
| Lysozyme | *LYZ* | Hs00426232_m1 |
| Mucin 2 | *MUC2* | Hs03005103_g1 |
| Neuronal differentiation 1 | *NEUROD1* | Hs01922995_s1 |
| Neurogenin 3 | *NEUROG3* | Hs01875204_s1 |
| Paired box 4 | *PAX4* | Hs00173014_m1 |
| Pancreatic and duodenal homeobox 1 | *PDX1* | Hs00236830_m1 |
| Peptide YY | *PYY* | Hs00373890_g1 |
| Somatostatin | *SST* | Hs00356144_m1 |

Supplementary Table 4. Reagents and Resources

| REAGENT or RESOURCE | SOURCE | CATALOG NUMBER |
|---|---|---|
| **Primary Antibodies (Dilution used)** | | |
| Alexa Fluor 647-Conjugated Anti-Chromogranin A Antibody (1:100) | Novus Biologicals | NBP2-47850AF647 |
| Alexa Fluor 647-Conjugated Mouse IgG2b kappa Isotype Control Antibody (1:100) | Biolegend | 400330 |
| Anti-Chromogranin A Antibody (1:100) | Agilent/Dako | M086901-2 |
| Anti-Chromogranin A Antibody (1:100) | Millipore Sigma | HPA017369-100UL |
| Anti-Cholecystokinin Antibody (1:100) | Abcam | Ab27441 |
| Anti-Cytokeratin 20 Antibody (1:50) | Thermo Fisher Scientific | 17329-1-AP |
| Anti-GIP Antibody (1:100) | Invitrogen | PA5-76867 |
| Anti-GLP-1 Antibody (1:100) | Abcam | Ab23468 |
| Anti-Lysozyme Antibody (1:50) | Novus Biologicals | NB100-63062 |
| Anti-MUC2 Antibody (1:50) | Novus Biologicals | NBP1-31231 |
| Anti-PYY Antibody (1:50) | Mybiosource | MBS9208739 |
| Anti-Serotonin Antibody (1:100) | Abcam | ab66047 |
| Anti-Somatostatin Antibody (1:100) | R&D Systems | mab2358 |
| APC anti-human β2-microglobulin Antibody (1:25) | Biolegend | 316311 |
| APC anti-human CD298 Antibody (1:25) | Biolegend | 341706 |
| PE-conjugated Anti-Chromogranin antibody (1:100) | BD Biosciences | 564563 |
| PE-conjugated Mouse IgG1 kappa Isotype Control Antibody (1:200) | BD Biosciences | 554680 |
| TotalSeq™-B0251 anti-human Hashtag 1 Antibody (1:25) | Biolegend | 394631 |
| TotalSeq™-B0252 anti-human Hashtag 2 Antibody (1:25) | Biolegend | 394633 |
| TotalSeq™-B0253 anti-human Hashtag 3 Antibody (1:25) | Biolegend | 394635 |
| TotalSeq™-B0254 anti-human Hashtag 4 Antibody (1:25) | Biolegend | 394637 |
| TotalSeq™-B0255 anti-human Hashtag 5 Antibody (1:25) | Biolegend | 394639 |
| TotalSeq™-B0256 anti-human Hashtag 6 Antibody (1:25) | Biolegend | 394641 |
| TotalSeq™-B0257 anti-human Hashtag 7 Antibody (1:25) | Biolegend | 394643 |
| TotalSeq™-B0258 anti-human Hashtag 8 Antibody (1:25) | Biolegend | 394645 |
| TotalSeq™-B0259 anti-human Hashtag 9 Antibody (1:25) | Biolegend | 394647 |
| **Secondary Antibodies** | | |
| Donkey anti-Goat IgG (H+L) Cross-Adsorbed Secondary Antibody, Alexa Fluor 488 (1:400) | Invitrogen | A-11055 |
| Donkey anti-Mouse IgG (H+L) Cross-Adsorbed Secondary Antibody, Alexa Fluor 488 (1:400) | Invitrogen | A-21202 |
| Donkey anti-Mouse IgG (H+L) Cross-Adsorbed Secondary Antibody, Alexa Fluor 647 (1:400) | Invitrogen | A-31571 |
| Donkey anti-Rabbit IgG (H+L) Cross-Adsorbed Secondary Antibody, Alexa Fluor 647 (1:400) | Invitrogen | A-31573 |
| Donkey anti-Rat IgG (H+L) Cross-Adsorbed Secondary Antibody, Alexa Fluor 488 (1:400) | Invitrogen | A-21208 |
| **Chemicals and Enzymes** | | |

| | | |
|---|---|---|
| 4′,6-diamidino-2-phenylindole (DAPI) | Thermo Fisher Scientific | D1306 |
| A-8301 | Millipore Sigma | SML0788 |
| AS1842856 | Millipore Sigma | 344355 |
| B-27 Supplement | Thermo Fisher Scientific | 12587010 |
| Betacellulin | Peprotech | 100-50 |
| Bovine serum albumin | Millipore Sigma | 05470-1G |
| Cell Recovery Solution | Corning | 354253 |
| Cell Staining Buffer | Biolegend | 420201 |
| Collagenase Type I | Thermo Fisher Scientific | 17018029 |
| DAPT | Selleckchem | S2215 |
| Diprotin A | Tocris | 6019 |
| Advanced DMEM/F12 | Thermo Fisher Scientific | 12634028 |
| DMEM Ca2+ Free | Thermo Fisher Scientific | 21068-028 |
| RNAse Inhibitor | Thermo Fisher Scientific | N8080119 |
| EGF, Recombinant Murine | Peprotech | 315-09 |
| Fetal Bovine Serum Certified USA Origin | Thermo Fisher Scientific | 16000044 |
| Forskolin | Tocris | 1099 |
| Gastrin I [Leu15], Human | Millipore Sigma | G9145 |
| GlutaMAX | Thermo Fisher Scientific | 35050061 |
| HEPES | Thermo Fisher Scientific | 15630080 |
| Matrigel, growth factor reduced, phenol red-free | Corning | 356231 |
| N-2 Supplement | Thermo Fisher Scientific | 17502001 |
| N-Acetyl-cysteine | Millipore Sigma | A7250 |
| Nicotinamide | Millipore Sigma | N0636 |
| Noggin, Recombinant Murine | Peprotech | 250-38 |
| Normocin | Invivogen | ant-nr-2 |
| Paraformaldehyde, 32% | Electron Microscopy Sciences (VWR) | 15714-S |
| PF-06260933 | Millipore Sigma | PZ0272 |
| Primocin | Invivogen | ant-pm-2 |
| Prolong Gold Antifade Mountant | Thermo Fisher Scientific | P36930 |
| Prostaglandin-E2 | Millipore Sigma | P0409 |
| Rimonabant | Millipore Sigma | SML0800 |
| SB202190 | Millipore Sigma | S7067 |
| SP600125 | Millipore Sigma | S5567 |
| Tranylcypromine | Tocris | 3852 |
| TRI Reagent® | Millipore Sigma | T9424-200ML |
| Triton X-100 | Millipore Sigma | T8787-50ML |
| TruStain FcX™ (Fc Receptor Blocking Solution), Human | Biolegend | 422301 |
| TrypLE Express | Thermo Fisher Scientific | 12605-010 |
| Tubastatin-A | Selleckchem | S8049 |
| Y-27632 dihydrochloride | Tocris | 1254 |
| **Commercial Assays** | | |
| 3’ Feature Barcode Kit, 16rxn Dual Indexing Compatible | 10X Genomics | 1000269 |

| | | |
|---|---|---|
| Click-IT EdU Alexa Fluor 488 Flow Cytometry Assay Kit | Thermo Fisher Scientific | C10420 |
| Chromium NextGem Chip G Single Cell Kit | 10X Genomics | 1000127 |
| Chromium Next Gem Single Cell 3’ GEM, Library and Gel Bead Kit v3.1, 4rxn Dual Index Compatible | 10X Genomics | 1000269 |
| Dead Cell Apoptosis Kit with Annexin V Alexa Fluor 488 & Propidium Iodide | Thermo Fisher Scientific | V13241 |
| Dead Cell Removal Kit | Miltenyi Biotec | 130-090-101 |
| Direct-zol RNA Microprep Kit | Zymo Research | R2061 |
| High-Capacity cDNA Reverse Transcription Kit | Thermo Fisher Scientific | 4368813 |
| Human GIP (total) ELISA Kit | Millipore Sigma | EZHGIP-54K |
| Human GLP-1 (7-36) ELISA Kit | Abcam | Ab184857 |
| Human PYY ELISA Kit | Abcam | Ab255727 |
| Human Serotonin ELISA Assay Kit | Eagle Biosciences/DLD Diagnostika GmbH | SER39-K01 |
| TaqMan™ Universal PCR Master Mix | Thermo Fisher Scientific | 4304437 |
